# Nanoparticle-mediated delivery of placental gene therapy via uterine artery catheterization in a pregnant rhesus macaque

**DOI:** 10.1101/2024.04.10.588902

**Authors:** Jenna K. Schmidt, Rebecca L. Wilson, Baylea N. Davenport, Timothy A. Hacker, Casey Fitz, Heather A. Simmons, Michele L. Schotzko, Thaddeus G. Golos, Helen N. Jones

## Abstract

Nanoparticles offer promise as a mechanism to non-invasively deliver targeted placental therapeutics. Our previous studies utilizing intraplacental administration demonstrate efficient nanoparticle uptake into placental trophoblast cells and overexpression of human *IGF1* (*hIGF1*). Nanoparticle-mediated placental overexpression of *hIGF1* in small animal models of placental insufficiency and fetal growth restriction improved nutrient transport and restored fetal growth. The objective of this pilot study was to extend these studies to the pregnant nonhuman primate and develop a method for local delivery of nanoparticles to the placenta via maternal blood flow from the uterine artery. Nanoparticles containing *hIGF1* plasmid driven by the placenta-specific *PLAC1* promoter were delivered to a mid-gestation pregnant rhesus macaque via a catheterization approach that is clinically used for uterine artery embolization. Maternal-fetal interface, fetal and maternal tissues were collected four days post-treatment to evaluate the efficacy of *hIGF1* treatment in the placenta. The uterine artery catheterization procedure and nanoparticle treatment was well tolerated by the dam and fetus through the four-day study period following catheterization. Nanoparticles were taken up by the placenta from maternal blood as plasmid-specific *hIGF1* expression was detected in multiple regions of the placenta via in situ hybridization and qPCR. The uterine artery catheterization approach enabled successful delivery of nanoparticles to maternal circulation in close proximity to the placenta with no concerns to maternal or fetal health in this short-term feasibility study. In the future, this delivery approach can be used for preclinical evaluation of the long-term safety and efficacy of nanoparticle-mediated placental therapies in a rhesus macaque model.

**Highlights:** Novel method to deliver therapeutics to maternal-fetal interface

Delivery of nanoparticles to the placenta via maternal catheterization

## Introduction

Placental health and the *in utero* environment are critical for maternal and fetal health. Aberrant placental development and function contribute to placental pathologies including fetal growth restriction (FGR), preeclampsia, premature rupture of membranes and preterm labor^1^. The impact of a poor *in utero* environment transcends pregnancy, predisposing the child to an increased risk of neurodevelopmental deficits and cardiovascular and metabolic disease in adulthood^2-5^. Current treatments for pregnancy complications are limited to premature delivery of the baby and the need for intensive neonatal care. There is thus a need to develop non-invasive placenta-targeted therapeutics to improve and extend placental health to term.

Nanoparticles offer promise as a clinically relevant vehicle for delivery of therapeutics to the placenta. We have previously demonstrated that a non-viral poly [2-hydroxypropyl] methacrylamide (HPMA) – poly(2-(N,N-dimethylamino) ethyl methacrylate (DMAEMA) co-polymer complexed with plasmid containing human insulin-like growth factor 1 (*hIGF1*) under the control of a placenta-specific promoter can be successfully delivered *in vivo* to pregnant mice^6^ and guinea pigs^7, 8^, *ex vivo* to human placental explants^9, 10^ and *in vitro* to human trophoblast cultures^6, 9, 10^. IGF signaling is essential to placental development and function by promoting trophoblast proliferation, syncytialization, invasion, migration and uptake of amino acids and glucose^11^. Lower levels of IGF1 have been observed in maternal circulation of human pregnancies complicated by FGR^12^, thus IGF1 is an excellent candidate for placental gene therapy. In our guinea pig maternal nutrient restriction model of FGR, nanoparticle-mediated *hIGF1* treatment increased expression of glucose and amino acid transport in the placenta, increased fetal glucose concentrations and stimulated downstream signaling to promote growth factor expression^7, 8^. Transgenic *hIGF1* was maintained for 5 days in the guinea pig placenta and multiple treatments beginning in mid-gestation positively impacted fetal growth. In human differentiated BeWo cells and placental explants studies, *hIGF1* nanoparticle treatment protected against cell death induced by oxidative stress and increased translocation of glucose transporter 1 to the cell surface membrane, respectively^10^. Altogether, nanoparticle-mediated *hIGF1* treatment in small animal models of placenta insufficiency and in cultured human trophoblasts has demonstrated improved placental function following treatment establishing the necessary mechanistic framework for transition into a nonhuman primate pregnancy model.

Nonhuman primates are translationally relevant models for human pregnancy as they share genetic features underlying embryonic and placental development and have a hemochorial placental structure that functions similarly to humans^13^. In a previous study, *hIGF*-containing nanoparticles were delivered by intraplacental injection around gestation day 100 and placenta and fetal tissues were analyzed at two- and ten-days post-treatment. Direct placental injection resulted in nanoparticle uptake by syncytiotrophoblasts as evident by detection of *hIGF1* transgene expression with corresponding downstream changes of proteins mediated by IGF signaling. The expression of the *IGF1* transgene, however, was restricted and focal to the injection site rather than being evenly distributed across cotyledons of both placental discs. The objective of the present study was to employ a uterine artery catheterization method, used clinically for uterine artery embolization, to deliver *hIGF1* containing nanoparticles into maternal circulation to non-invasively maximize the opportunity for uptake by trophoblasts, and thus uniformly treat the placenta.

## Methods

### Nanoparticles

Detailed methods for synthesis of the HPMA_115_-DMAEMA_115_ co-polymer and the complexing of the plasmid to form the nanoparticle has been previously described^8^. The plasmid coding for human *IGF1* driven by a trophoblast-specific promoter, *PLAC1*, was produced using previously reported methods^8^.

### Delivery of nanoparticles via uterine artery catheterization

The pregnant rhesus macaque (*Macaca mulatta*) enrolled in this study was 5.1 years old and weighed 5.9 kg. The animal was monitored at least twice daily for general health and well-being. The female was co-housed with a compatible male upon observation of menses and pregnancy was confirmed by abdominal ultrasound. The gestational age was estimated by measuring fetal crown rump length. The dam was anesthetized with an intramuscular dose of ketamine (10 mg/kg) for ultrasound examination of pregnancy health and blood draws.

The uterine artery catheterization was performed on gestation day 108 at the University of Wisconsin-Madison Cardiovascular Physiology Core Facility. The dam was sedated with ketamine (intramuscular dose of up to 20 mg/kg), midazolam (0.1-0.3 mg/kg IM), and propofol (1-5 mg/kg IV). General anesthesia was maintained throughout the course of the procedure using a combination of isoflurane gas (0.25-4% isoflurane/O_2_) by way of an endotracheal tube and a constant rate of fentanyl (2-23 mcg/kg/h IV) or lidocaine (up to 2 mg/kg) and buprenorphine (up to 0.005 mg/kg) combined were administered to the epidural space. Isotonic intravenous fluids were administered at a rate of 1-10 ml/kg/h and as a bolus to counteract blood flow pressure and buprenorphine SR (0.2 mg/kg SQ) was administered. The dam was monitored using a pulse oximeter, capnograph, blood pressure monitor, and temperature probe. The female was placed in the supine position. After a sterile preparation of the skin near the femoral artery, a 5Fr sheath (Terumo) was placed in the femoral artery percutaneously. An 0.035 guidewire was placed in the sheath and directed to contralateral uterine artery under fluoroscopic guidance. A 5Fr catheter (Cobra C2, Merit) was placed over the wire to the uterine artery. Placement of the catheter was by infusion of a 1-2ml bolus of contrast agent (iohexol, 50 mgl/ml). Heparin was given as an anticoagulant at a dose of 1000 units. A volume of 5.0 mL containing 500 μg of nanoparticles was infused into the catheter and followed by sterile saline to flush the catheter. Pregnancy health was evaluated by ultrasound examination of fetal heart rate prior to and following the uterine artery catheterization procedure.

### Blood and Tissue Collection

A blood sample was drawn 7 days prior to the delivery of nanoparticles, prior to uterine artery catheterization delivery of nanoparticles, and then 4 h, 2 days and 4 days post-nanoparticle delivery. Blood was drawn from the femoral or saphenous vein using a vacutainer system. Blood tubes were centrifuged at 1300 x *g* for 10 min at room temp and the serum and plasma were aliquoted and frozen at −80ºC. Complete blood count tests were performed on the fresh blood sample as previously described^14^.

At 4 days post-treatment with nanoparticles, an ultrasound was performed to confirm fetal viability prior to the removal of the fetoplacental unit via hysterotomy. Anesthesia was performed as similarly described above for the uterine artery catheterization procedure. During the hysterotomy procedure, the entire conceptus (i.e., fetus, placenta, fetal membranes, umbilical cord, and amniotic fluid) was removed. The fetus was euthanized by intracardiac injection of 50 mg/kg sodium pentobarbital. Biopsies of maternal uterine-placental bed, mesenteric lymph node, liver, and spleen were collected aseptically and the dam was recovered. The dam received bupivacaine liposome injectable suspension (Nocita, up to 10.6 mg/kg, SQ or IM) injected during body wall closure as a nerve block and a combination of buprenorphine (0.01-0.03 mg/kg IM), buprenorphine SR (0.2 mg/kg SQ), and meloxicam (0.1-0.2 mg/kg SQ or PO) as the post-op analgesia regimen.

Maternal-fetal-interface and fetal tissues were processed for histology, in situ hybridization and RNA isolation. A center cut of the placenta containing the umbilicus was collected for histopathological analysis. Each placental disc was then dissected into 6 regions using a grid pattern and from each grid piece a sample of tissue was preserved by flash freezing in liquid nitrogen for *in situ* hybridization, collected into RNAlater stabilization solution (ThermoFisher Scientific, cat no. AM7021), or placed into a cassette for fixation and subsequent histopathological analysis. Archived placental flash-frozen tissue and RNA samples from gestationally age-matched control pregnancies (n=6) were obtained from Dr. Golos to compare to the nanoparticle treated samples.

### Hormone Assays

Steroid hormones in serum were assessed by the Wisconsin National Primate Research Center’s Assay Services Unit using a Roche Cobas e411 analyzer as previously described^15^.

### Histopathology

Tissues were fixed in 4% PFA for 24 h, transferred to 70% ethanol, processed in an automated tissue processor and embedded in paraffin. Paraffin embedding, tissue sectioning and hematoxylin and eosin (H&E) staining were performed by the Comparative Pathology Lab through the University of Wisconsin Research Animal Resources and Compliance laboratory. Histological evaluation of tissues was performed by a board-certified veterinary pathologist.

### In situ hybridization

In situ hybridization (ISH) was performed as previously described using a custom designed BaseScopeTM probe (Advanced Cell Diagnostics) to detect plasmid-specific *IGF1* expression at 96 h after nanoparticle delivery^8^.

### RNA extraction and Quantitative PCR (qPCR)

Gene expression analysis was performed as previously described^8^. RNA was extracted from 50 mg of snap frozen placenta tissue using the RNAeasy Mini kit (Qiagen) following standard manufacturers protocol and including a DNase treatment. A High-capacity cDNA Reverse Transcription kit (Applied Biosystems) was used to convert 1 μg of RNA to cDNA. The qPCR reactions were assembled using a PowerUp SYBR green kit (Applied Biosystems) and predesigned primers (Sigma KiQStart Primers, β-ACTIN, TBP and GAPDH). Human-specific *IGF1* primers were designed for species specificity *(hIGF1; Fwd: 5’-* CGCTGTGCCTGCTCACCT, Rev: 5’ – TCTCCAGCCTCCTTAGATCA) and purchased from Integrated DNA Technologies (IDT). Specificity was tested using RT-PCR (Sigma Fast Start PCR mastermix) on both gDNA and cDNA for 35 cycles using both human and rhesus macaque samples. Amplified samples were then run on a 1.5% agarose gel and imaged using a BioRad ChemiDoc to confirm species specificity before performing qPCR experiments. The qPCR reactions were performed using the Quant3 Real-Time PCR System (Applied Biosystems). Gene expression was normalized using reference genes β-ACTIN, TBP and GAPDH and expression was calculated using the comparative CT method with the Design and Analysis Software v2.6.0 (Applied Biosystems).

## Results

### Delivery of nanoparticles via uterine artery catheterization

To deliver nanoparticles into maternal circulation, the uterine artery was accessed percutaneously and a 5Fr sheath placed to facilitate the use of a 0.35 guidewire with a slight curve placed on the end to direct the wire to the contralateral iliac where the aorta bifurcates to the iliac arteries typically around the fourth lumbar vertebra. The catheter was used to support the wire and then once the wire was inserted into the contralateral femoral artery for extra support, the catheter was advanced to the contralateral iliac. Once the catheter was in the iliac, the wire was pulled back to internal iliac to access the uterine artery. The catheter could be advanced at this point to the uterine artery. The fluoroscopic camera was perpendicular to the animal positioned on its back with legs spread wide. A small amount of contrast was used to identify the uterine artery as shown in Figure 1. The main challenge is driving the wire and catheter to the contralateral iliac artery. A gentle curve on your wire and flexible catheter with a slight bend will help to direct your hardware to the uterine artery. A volume of 5.0 ml of nanoparticles containing plasmid coding for *hIGF1* driven by the trophoblast specific *PLAC1* promoter was injected into maternal circulation for direct perfusion of nanoparticles into the placenta enabling contact and uptake by the syncytium.

**Figure 1.**
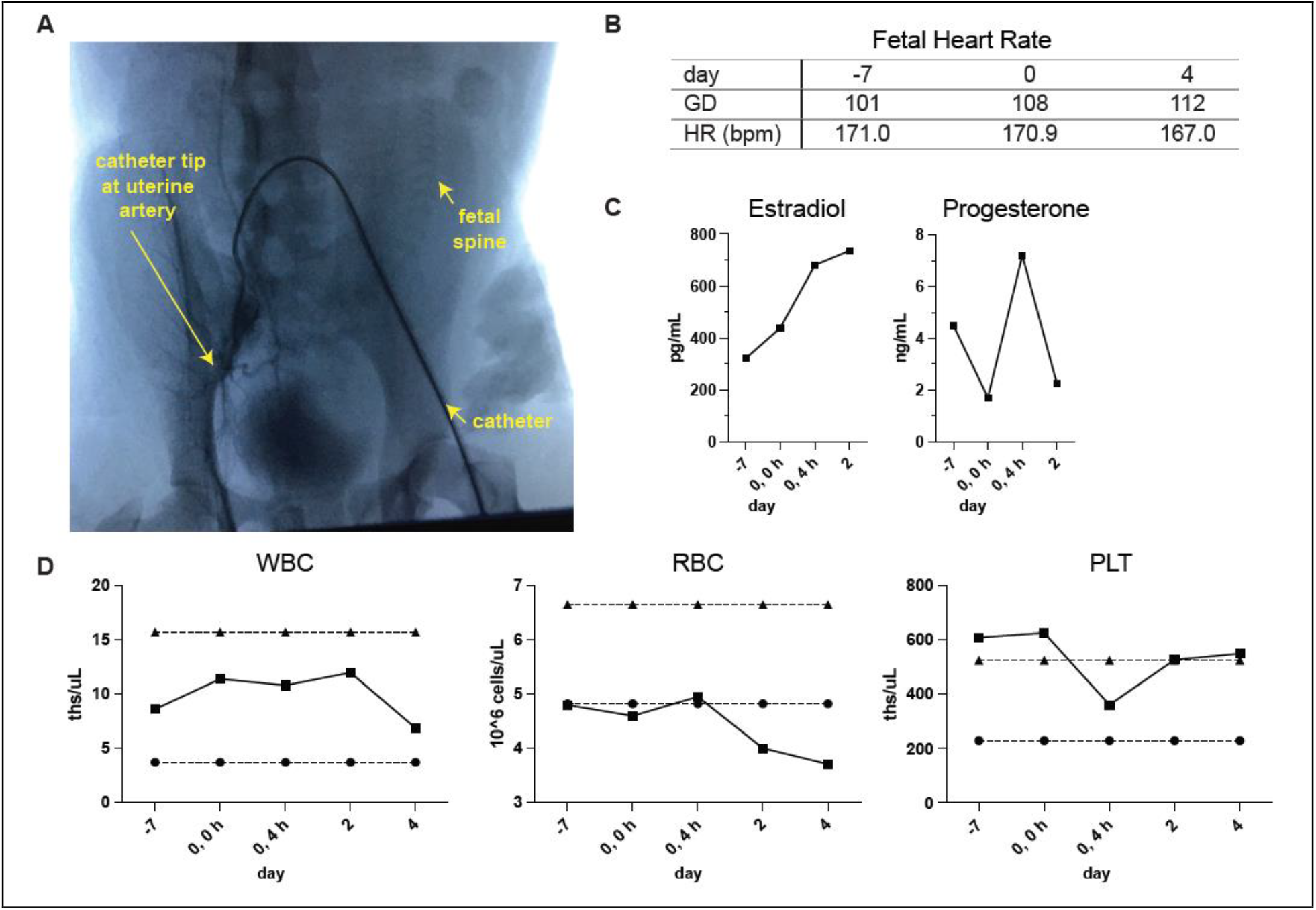
Evaluation of acute maternal and fetal responses to uterine artery catheterization. A. Image of fluoroscopy-guided catheterization of the uterine artery. The arrows indicate the catheter placement in the femoral artery, the tip of the catheter at the uterine artery, and the fetal spine relative to the uterine artery. B. Fetal heart rate (HR) in beats per minute (bpm) at −7 days prior to treatment, at the time of the procedure and 4 days post-treatment. C. Levels of estradiol and progesterone in maternal circulation. D. Complete blood count analysis of maternal white blood cells, red blood cells, and platelet levels as evaluated by complete blood cell count. The reference interval for each parameter is indicated by dashed lines.

The dam and fetus were monitored closely following nanoparticle delivery. Fetal heart rate measures were within normal range by ultrasound when examined prior to treatment, immediately following treatment and at 4 days post-treatment^16^. Fetal growth measures by ultrasound were also within range for the gestational age at the time of treatment (biparietal diameter 3.65 cm, femur length 2.50 cm)^16^. Estradiol and progesterone levels in maternal circulation varied (Figure 1C) and were within normal range for gestational age (unpublished data). A complete blood count on maternal blood was performed prior to treatment and at 4 h, 2 days and 4 days post-treatment revealing that her white blood cell (WBC) counts were within expected ranges. The red blood cell (RBC) count tended to decline with additional blood draws post-treatment, however, the mean RBC volume and mean hemoglobin amount per RBC were relatively unchanged and within normal reference intervals. The number of platelets declined after the procedure and then rebounded by day 2 post-treatment.

The pregnancy was terminated at 4 days post-treatment to evaluate nanoparticle-mediated delivery of the *hIGF1* transgene. The female fetus and placenta were grossly unremarkable and tissue weights were within normal ranges for gestational age (fetus 191.40 g, total placenta 71.11 g, placenta disc 1 48.22 g, placenta disc 2 22.89 g)^17^. There were no significant histologic lesions in maternal tissue biopsies of liver, spleen and lymph node nor in fetal tissues including brain, liver, heart, spleen, kidney, and lung.

### Expression of hIGF1 transgene

To evaluate nanoparticle-mediated gene therapy, ISH and qPCR were performed using probes and primers that distinguish between plasmid-specific *hIGF1* expression from endogenous macaque *IGF1*. At four days post-treatment, *hIGF1* signal was still detectable and present in all six regions of the placenta that were evaluated (Figure 2A,B). *hIGF1* expression was detected near vessels within the decidua (Figure 2A, C, E-F) and in the chorionic plate (Figure 2D). Plasmid-specific *hIGF1* expression was further confirmed by qPCR with elevated expression in the nanoparticle-treated placenta and undetectable in untreated control placenta (Figure 2C).

**Figure 2.**
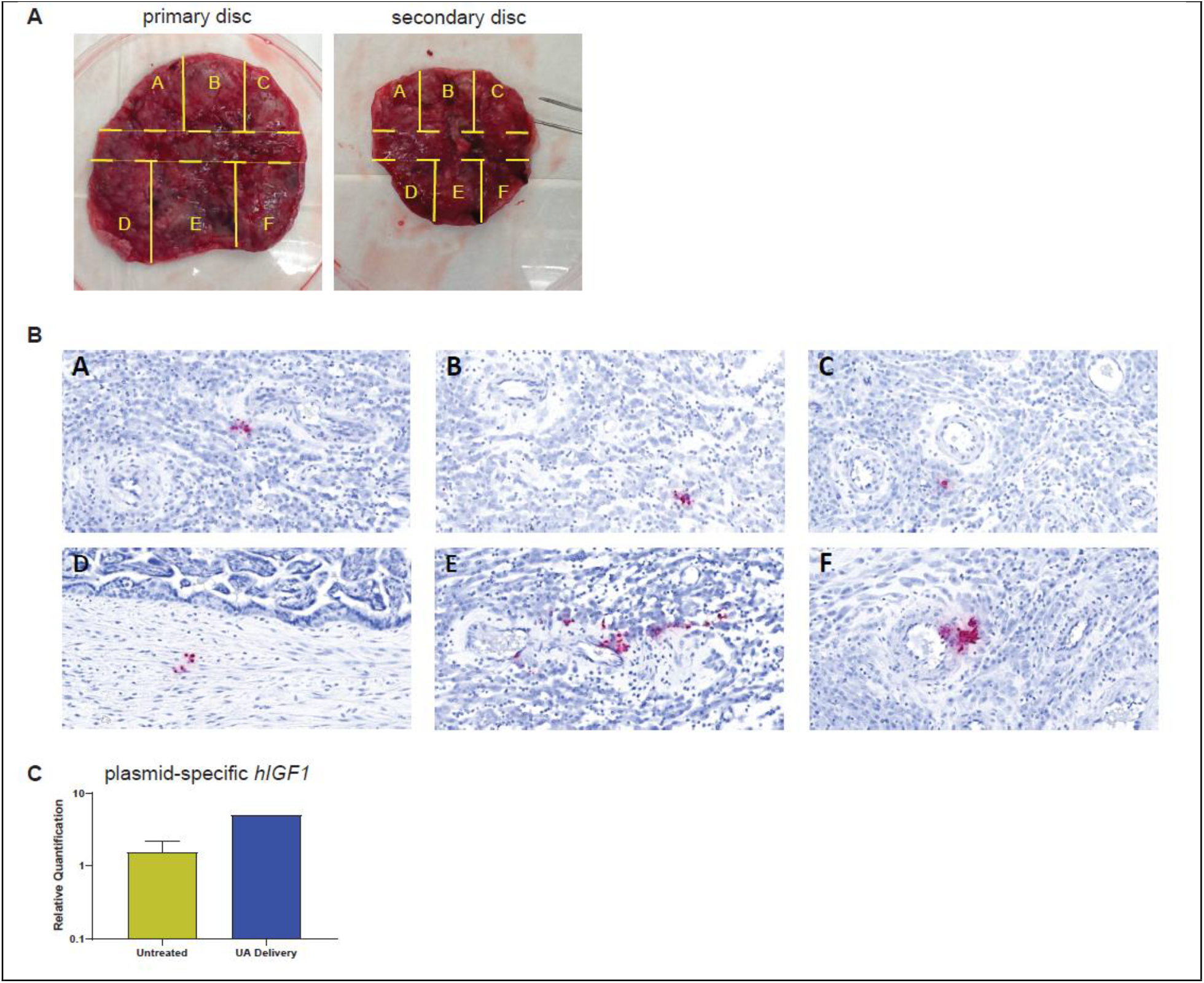
Detection of *hIGF1* in the nanoparticle-treated placenta. A. Gross images of the placental discs. Each disc was dissected into six regions using a grid pattern to evaluate RNA and protein expression. A center cut (tissue region indicated by the dashed lines) was collected for histopathological analysis. B. In situ hybridization of tissues pieces A-F of the primary disc. C. Relative gene expression of *hIGF1* in control, untreated placenta compared to the placenta that received nanoparticles via uterine artery (UA) delivery as determined by qPCR.

## Discussion

Pregnancy complications stem from abnormal placental development and function, yet treatments to improve placental health and maintain pregnancy remain limited. While intraplacental injection of a treatment is not a clinically relevant delivery method, the catheterization of the uterine artery is clinically relevant (i.e., used for embolization of uterine fibroids) and allows for less invasive delivery to the maternal-fetal interface. In this proof-of-concept study, we showed successful delivery of nanoparticles into the uterine artery of a pregnant rhesus macaque using fluoroscopy to guide a catheter from the femoral artery, through the common iliac artery, around the bend in the abdominal aorta, down the contralateral common iliac and then advanced into the uterine artery. The procedure was well-tolerated by the dam and fetus with observations of maternal and fetal health being unremarkable through the short duration of study.

Nanoparticles containing *hIGF1* plasmid in our previous guinea pig and nonhuman primate studies were directly injected into the placenta under ultrasound guidance. In the guinea pig studies, *hIGF1* signal remained after five days of treatment and treatment increased placental expression of glucose transporter, *Slc2A3* with corresponding increases in glucose concentration in the fetuses^8^. Translating these studies into macaques which more closely model human pregnancy, we previously showed successful *hIGF1* transgene expression at two and ten days post-intraplacental injection (Wilson et al., under review). The *hIGF1* signal in the macaque syncytiotrophoblasts was focal, and likely restricted to the injection site at four days post-treatment. In the current study, the delivery of nanoparticles into maternal circulation enhanced the opportunity for broader uptake across the placenta. Indeed, *hIGF-1* signal was still detectable at four days post-delivery into the uterine artery and our preliminary analysis suggests treatment stimulates *IGF1* signaling pathways.

The observations of this report are presented as a case study of a single animal. Further studies are needed in a larger cohort of macaques to thoroughly evaluate the efficacy and safety of nanoparticle-mediated *hIGF1* treatment delivered via uterine artery catheterization on maternal and fetal health. Additional studies are also needed to assess the safety of multiple delivery events during pregnancy, as the ultimate goal is to sustain *IGF1* signaling throughout the third trimester until term delivery in human pregnancies complicated by FGR. Overall, this study establishes a method to deliver treatments directly into maternal blood that perfuses the placenta.

## Abbreviations

(FGR): fetal growth restriction
(hIGF1): human insulin-like growth factor 1

## Ethics Statement

The animal was cared for by the staff at the Wisconsin National Primate Research Center and all procedures were performed in accordance with the NIH Guide for the Care and Use of Laboratory Animals and under the approval of the University of Wisconsin College of Letters and Sciences and Vice Chancellor Office for Research and Graduate Education Institutional Animal Care and Use Committee (protocol G006040).

## Acknowledgments

We would like to thank the Wisconsin National Primate Research Center’s Veterinary Services, Scientific Protocol Implementation, Pathology Services, Assay Services and Animal Services Unit staff for providing animal care, and assisting in procedures including breeding, pregnancy monitoring, anesthesia support, hematology, and sample collection.

## Author Contributions

RLW performed experiments, analyzed data and wrote manuscript. JKS designed, oversaw experiments and contributed to writing of the manuscript. BND performed experiments and analyzed data. TH developed and performed the uterine artery catheterization procedure. CF aided in the catheterization procedure and provided surgical support. HAS collected tissues and performed histopathological analysis. MLS performed animal procedures and collected samples. TGG designed and oversaw experiments. HNJ obtained funding, conceived the study and edited the manuscript. All authors have approved the final version.

